# Probable omnigenic effect and evolutionary insights of aerobic adaptation allele OsNCED2^T^ on qDTY12.1 effecting grain yield under reproductive stage drought stress in rice

**DOI:** 10.1101/2021.03.10.434074

**Authors:** C Parameswaran, B Cayalvizhi, S Sanghamitra, N Anandan, K Jawahar Lal, BN Devanna, K Awadhesh, J Kishor, S Bandita, B Niranjana, B Biswaranjan

## Abstract

Yield associated quantitative trait loci (qDTY) under drought stress provides significant advantage for grain yield in rice. The major, stable qDTY12.1 was identified in a mapping population developed from upland cultivars Vandana and Way Rarem. Further, introgression line comprising of qDTY12.1 genomic region was characterized to have multiple genes (NAM, DECUSSATE) regulating the drought tolerance under severe drought stress substantiated through recently proposed omnigenic model for complex traits. Recently, plastid localized NCED2^T^ allele present within the qDTY12.1 genomic region was characterized for conferring aerobic adaptation in lowland varieties. Since, NCED2^T^ is evolutionary fixed in upland cultivars and Vandana was found to have the favorable allele of NCED2^T^, we hypothesized that this favorable allele might confer omnigenic effect on qDTY12.1 genes. Our evolutionary analysis using non synonymous SNPs present in genes namely NCED, NAM, and DECUSSATE and qDTY12.1 genomic regions showed specific grouping of Vandana with upland cultivars only for NCED gene and its adjoining genomic regions. However, non synonymous SNPs in NAM and DECUSSATE genes and its adjoining genomic regions of drought tolerant varieties were closely related and grouped together in the phylogenetic analysis. Moreover, ecotype specific differentiation and greater nucleotide difference with wild relatives was also observed for DECUSSATE gene in rice. This finding indicates differential evolution of qDTY12.1 regions for upland and drought tolerance and omnigenic effect of NCED2^T^ gene in qDTY12.1. Further, we propose a breeding model for enhancing genetic gain for yield under severe drought stress by incorporation of NCED^T^, qDTY12.1 and other drought tolerant QTLs for membrane stability in rice.

## Introduction

Abscisic acid (ABA) is a major plant hormone which plays a significant role in drought tolerance in plants (Zhang et al, 2006). Plants adaptation to water limitation conditions are tightly regulated through abscisic acid biosynthesis (Xiong and Zhu, 2003). The plastid localized NCED genes (9-*cis*-epoxy-carotenoid dioxygenase) are involved in biosynthesis of precursors required for the synthesis of ABA in plants (Vallabhaneni et al, 2010). In rice, favorable allele of NCED2^T^ was specifically fixed in upland cultivars and reported to increase the cellular ABA levels and lateral roots (Lyu et al, 2013). Recently, NCED2^T^ allele was functionally characterized for the aerobic adaptation in upland cultivars (Hu et al, 2020). This gene is located in the chromosome number 12 of rice at 14, 232,905 to 14,234,854 Mbp.

Genetic studies on drought tolerance for yield in rice resulted in the identification of several yield QTLs (qDTY) which relatively maintains yield under drought stress conditions (Kumar et al, 2014). A stable, major, large effect QTL for drought tolerance namely qDTY12.1 enhanced drought tolerance of lowland cultivars (Mishra et al, 2013). This QTL was identified and fine mapped using Vandana and Way Rarem mapping population and located within the RM28048 (14,106,460) and RM28166 (17,607,668) markers located in chromosome 12. Besides, peak marker (RM512) was found to be present at 17,395,485 bp in chromosome 12 of this QTL (Dixit et al, 2012). Further characterization of qDTY12.1 showed multiple genes within the QTL region might be regulating the drought tolerance response for yield in rice (Dixit et al, 2015). Specifically, two genes namely *OsNAM*_*12*.1_: 17,391,342 to 17,392,627 (Dixit et al, 2015) and *OsDEC*: 16,500,219 to 16,504,887 were characterized through multiple approaches as casual genes providing ‘omnigenic effect’ for drought tolerance in the qDTY12.1 (de los Reyes et al, 2021).

Furthermore, introgression line used in the fine mapping and characterization of qDTY12.1 spanned between RM28048 and RM28166 of qDTY12.1. In addition, both the parents namely Vandana and Way Rarem were upland cultivars. Besides, our preliminary observation using SNP seek database also showed presence of OsNCED2^T^ allele in Vandana cultivar. Therefore, we hypothesized that both the parents might have OsNCED2^T^ allele and this favorable allele responsible for efficient ABA synthesis might provide ‘omnigenic effect’ to the causal genes present in qDTY12.1.

## Methodology

SNP seek database of rice was used to retrieve the non synonymous single nucleotide polymorphisms in the selected genes and within qDTY12.1 (Alexandrov et al, 2015). Phyogeny.fr online tool was used for construction of phylogenetic analysis using default parameters (Dereeper et al, 2008). TASUKE tool was used for retrieval of gene sequence from wild relatives of rice (Kumagai et al, 2013). MEGAX software was used for the multiple alignment and neighbor-joining phylogenetic analysis of retrieved sequences (Kuamr et al, 2018). Flapjack tool was used for the graphical representation of the non synonymous SNPs in the genes (Milne et al, 2010)

## Results

The presence of OsNCED2^T^ allele (LOC_Os12g24800) in 3k rice panel was initially analyzed in SNP seek database of rice. The allelic position (14233796) corresponding to the functional alternate alleles (C/T) in the NCED2 gene showed approximately 9% (268 nos) of rice genotypes in 3k panel is having the favorable OsNCED2^T^ allele. Besides, heterozygous alleles were found in 75 rice genotypes. Further, *indica 3, indx*, and *sub tropical japonica* has more number of genotypes containing favorable allele as compared to other ecotypes (Fig.1). Apart from the functional allele, all non-synonymous SNPs identified against Nipponbare reference of the three selected genes (NCED, NAM, DEC) were also analyzed. These three genes were previously characterized for drought tolerance and present within qDTY12.1. Non synnonumous SNPs for the three genes were retrieved from seven selected genotypes namely N22 (drought tolerant landrace), Vandana (upland variety), IRAT 104 (upland variety), Swarna, Bala, and FL478 from SNP seek database. The analysis identified 14, 3, and 22 SNPs in NCED, NAM, DEC genes, respectively. Further, phylogenetic reconstruction of selected genotypes was performed using the non-synonymous SNPs identified in the three genes (Fig. 2).

**Fig. 1.**
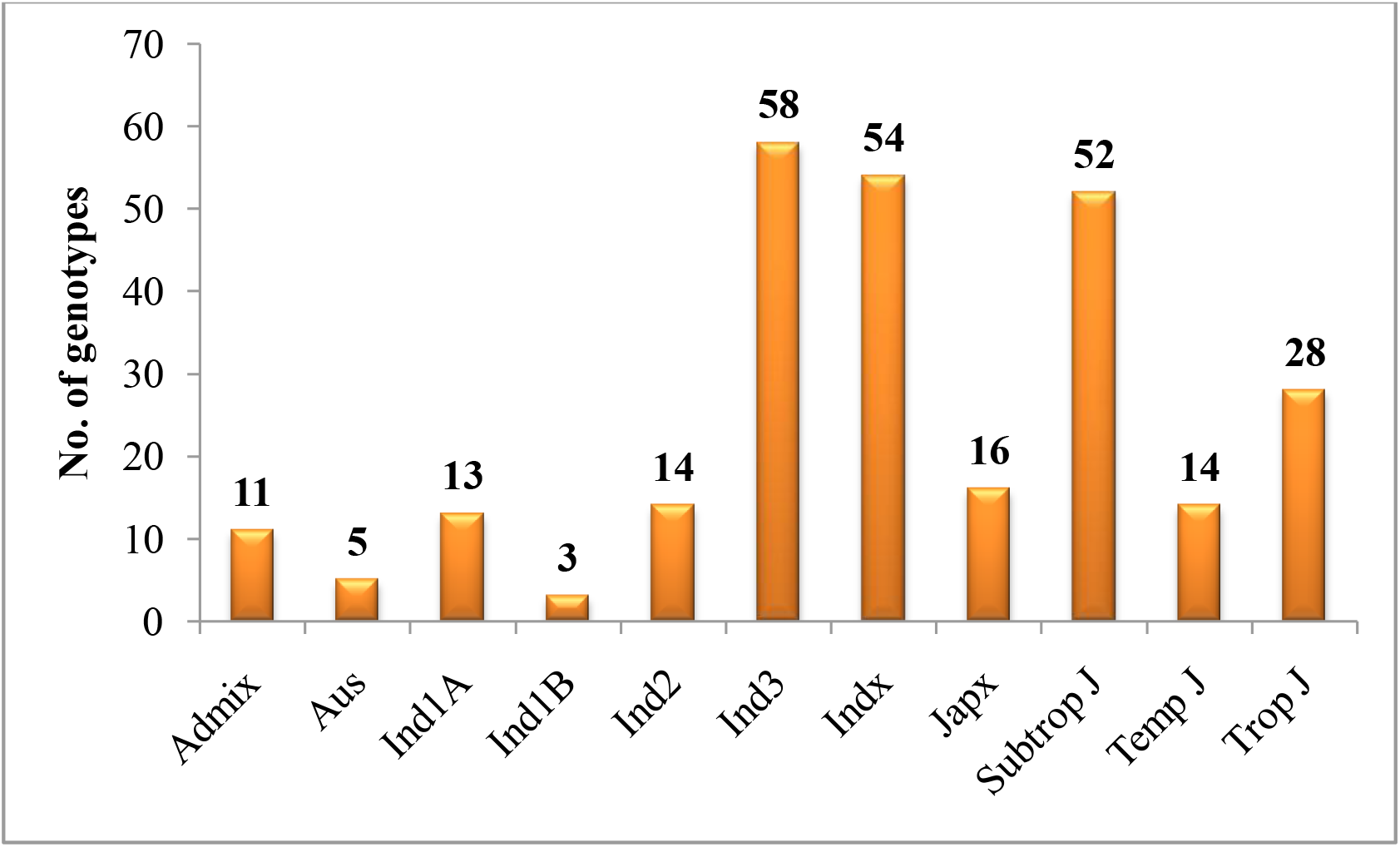
Distribution of favorable allele (OsNCED2^T^) in different rice accessions in SNP seek database

**Fig. 2.**
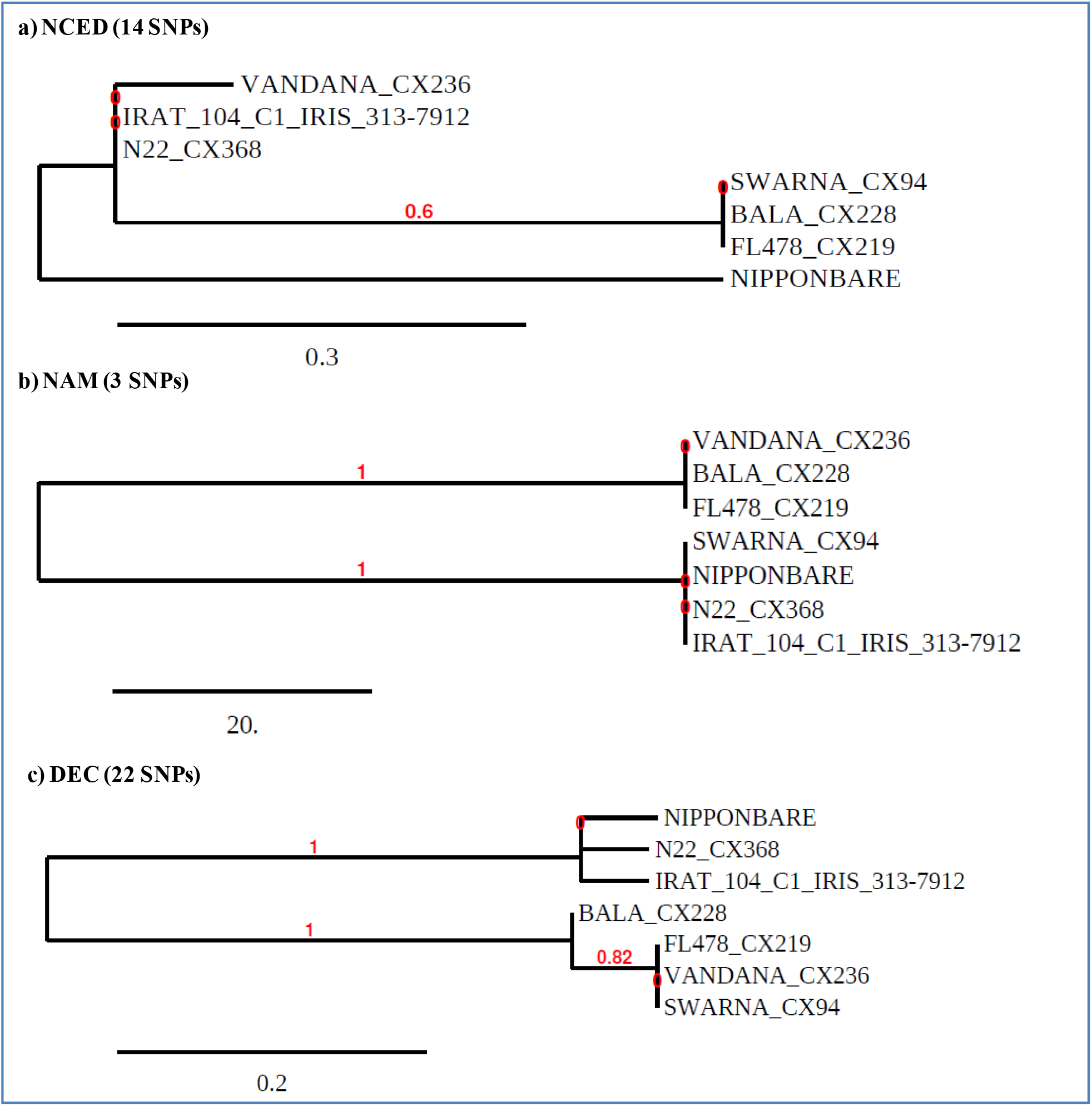
Phylogenetic analysis of seven genotypes using SNPs present in three selected genes. a-neighbor-joining phylogeny using non synonymous SNPs in NCED gene, b-neighbor-joining phylogeny using non synonymous SNPs in NAM gene, c-neighbor-joining phylogeny using non synonymous SNPs in DECUSSATE gene. Numbers in bracket indicates total non synonymous SNPs identified in the three selected genes.

The phylogeny for NCED gene showed seven genotypes were separated into two clusters and Nipponbare was present as outgroup. The first cluster consists of drought tolerant and upland varieties namely N22, Vandana, and IRAT 10. The remaining susceptible genotypes were present in cluster II of the phylogeny. Similarly, phylogeny of NAM gene divided seven genotypes into two clusters. The upland or drought tolerant cultivar namely N22, IRAT 104 were present in one group along with Swarna and Nipponbare. However, Vandana, Bala, and FL478 were present in cluster II. Also, N22 and IRAT 104 were present in same cluster for DEC gene in the phylogeny. Thus, initial phylogenetic analysis for the functional genes clearly showed drought tolerant/upland cultivar, N22 and IRAT 104 were always grouped in same cluster for three genes. In contrast, Vandana was grouped along with N22 and IRAT 104 only for NCED gene and were grouped together with susceptible cultivars for remaining two genes.

We hypothesized that differential evolution of genomic regions within the qDTY12.1 might be the reason for the grouping of upland genotype Vandana with other genotypes namely N22 and IRAT 104. Hence, phylogenetic reconstruction was again performed using all the non synonymous SNPs spanning the qDTY12.1 for the seven genotypes. The qDTY12.1 (14-18 Mbp) in chromosome 12 were divided into four different genomic blocks (14-15 Mbp, 15-16 Mbp, 16-17 Mbp, 17-18 Mbp) and analyzed for the grouping of selected genotypes. The analysis showed there were 681, 322, 306, and 180 non synonymous SNPs within the four genomic bocks, respectively. Further, Vandana, N22, and IRAT 104 were grouped together only in 14-15 Mbp genomic region of qDTY12.1. But, Vandana was grouped along with the susceptible genotypes and N22 and IRAT 104 were present in same cluster for the 15-16 Mbp genomic region of qDTY12.1. These findings showed not only the selected genes but there was also differential evolution of genomic regions within qDTY12.1. Accordingly, we have predicted the genomic regions for the genotypes N22, Vandana, IRAT 104, and Way Rarem (Fig. 3f). The entire qDTY12.1 was divided into two genomic segments or blocks. The first segment comprises of favorable allele of NCED2^T^ present only in upland genotypes namely Vandana, 1RAT 104, and Way Rarem. The second block comprises of genomic regions specifically in N22, IRAT 104 and Way Rarem. Further, first and second genomic block in qDTY12.1 provides early drought response and yield maintenance under drought, respectively. Thus, we assume that NCED2^T^ allele might have an omnigenic effect on the genomic blocks of qDTY12.1.

**Fig. 3.**
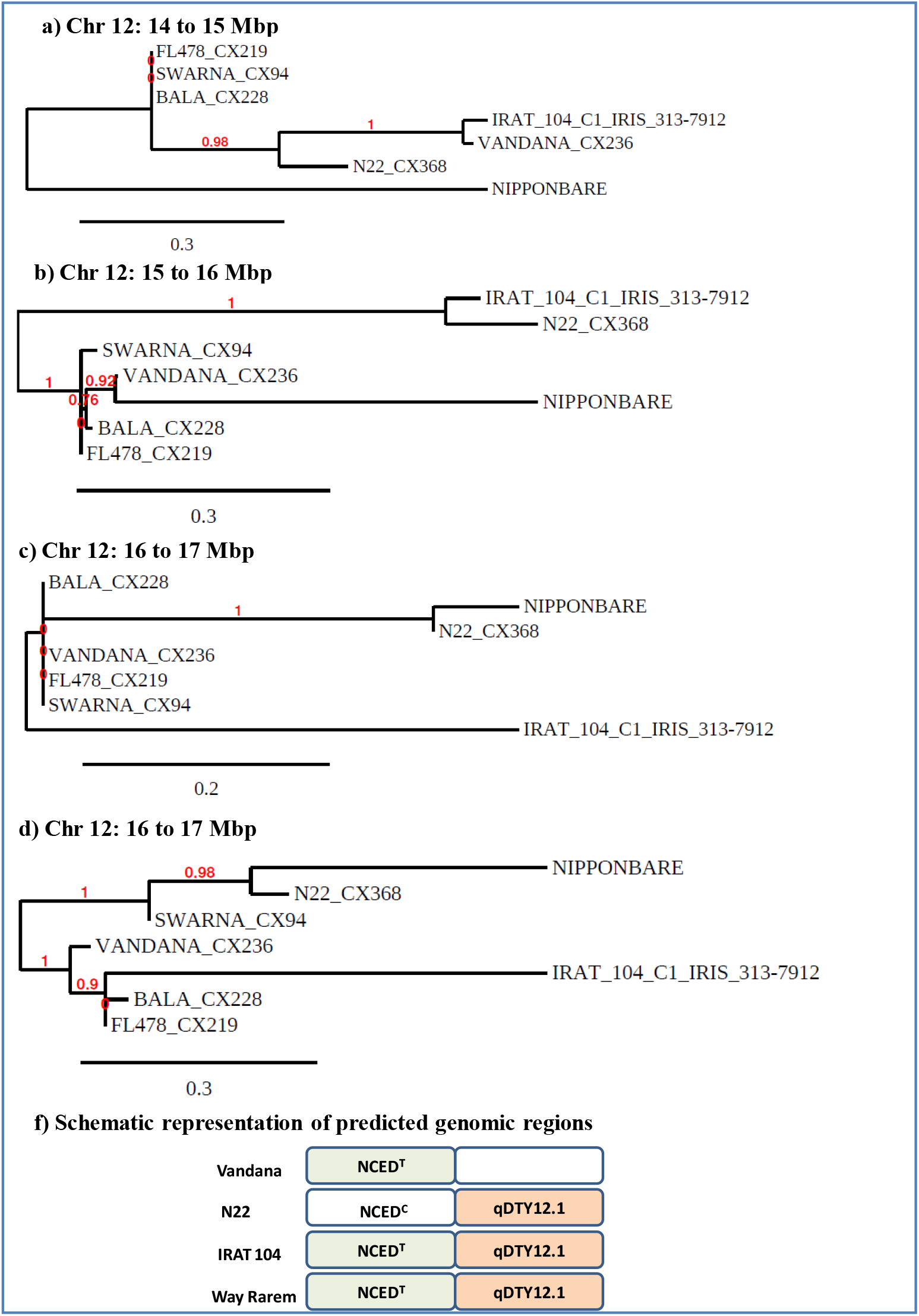
Phylogenetic analysis of seven genotypes using SNPs identified in qDTY12.1. a,b,c,d represents neighbor-joining phylogenetic tree for different genomic regions. f-represents two genomic blocks in qDTY12.1 and predicted genotypes for four cultivars. Green color fill indicates presence of favorable allele for NCED gene, brown color fill indicates favorable alleles for qDTY 12.1

The presence of the favorable allele in OsNCED2^T^ was also analyzed in wild relatives of rice. The analysis showed only one accession of *Oryza rufipogan* (W1807) has the favorable allele for NCED2 gene (Fig. 4). Then, non synonymous substitutions in the DECUSSATE gene were also analyzed in all rice genotypes of the 3k panel and wild relatives. The haplotype blocks was constructed in different rice ecotypes for the DECUSSATE gene and analyzed for the specific fixation of non reference alleles (Fig. 5). The analysis showed sub tropical and temperate japonica retained the reference alleles of Nipponbare. However, all other ecotypes including tropical japonica showed non synonymous substitutions in the gene of varying proportions. Additionally, many wild relatives also showed non synonymous substitutions in the coding regions of the gene (Fig. 6). Thus, DECUSSATE gene and the genomic regions of the qDTY12.1 might have evolutionary significance in rice.

**Fig. 4.**
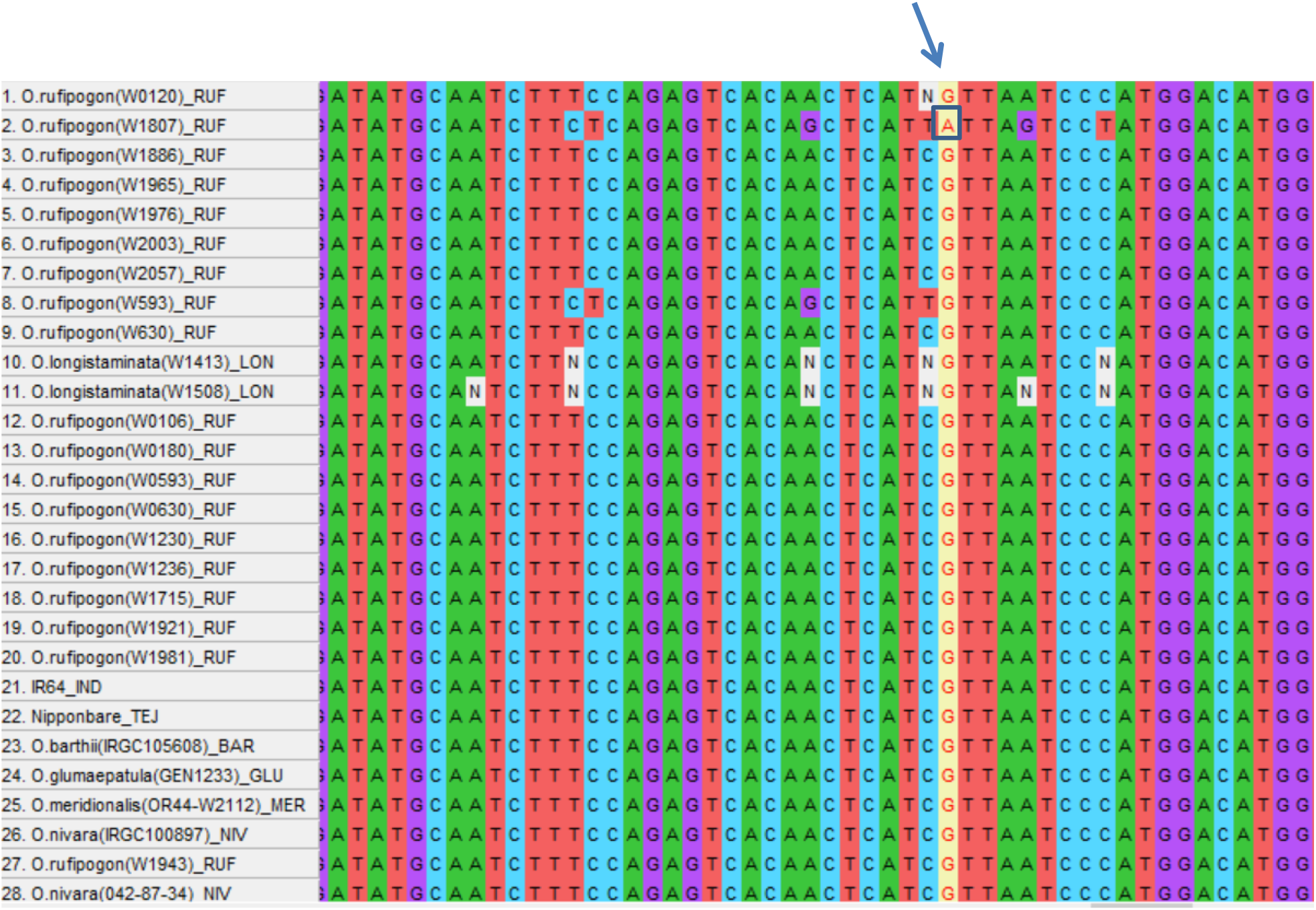
Multiple alignment of NCED gene in rice wild relatives. Arrow indicates the alignment position and boxed nucleotide indicate the favorable allele in one of the wild relatives (*Oryza rufipogan*).

**Fig. 5.**
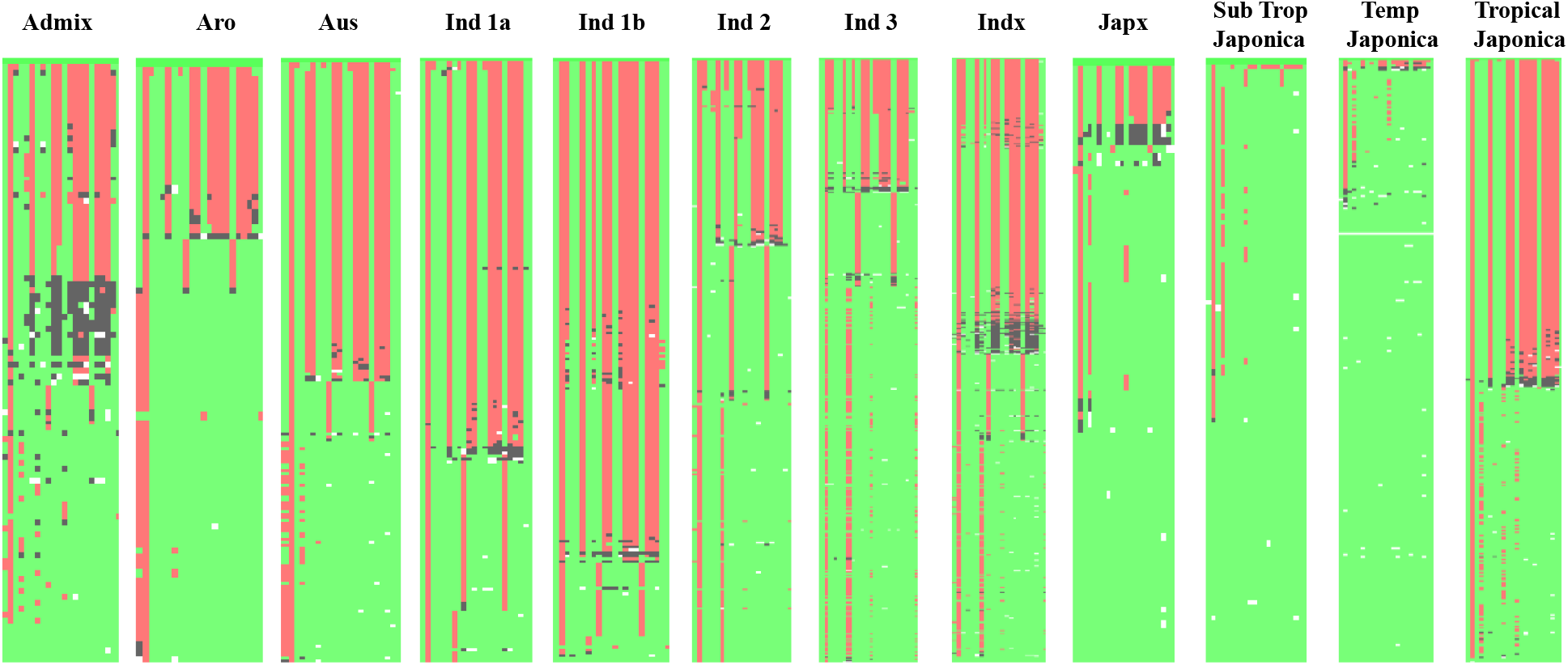
Graphical representation of haplotypes of DECUSSATE gene in different ecotypes of rice Green, red and black represents similar, alternate and heterozygous allele to Nipponbare reference gene sequence. Non availability of sequence information was indicated in white color boxes.

**Fig. 6.**
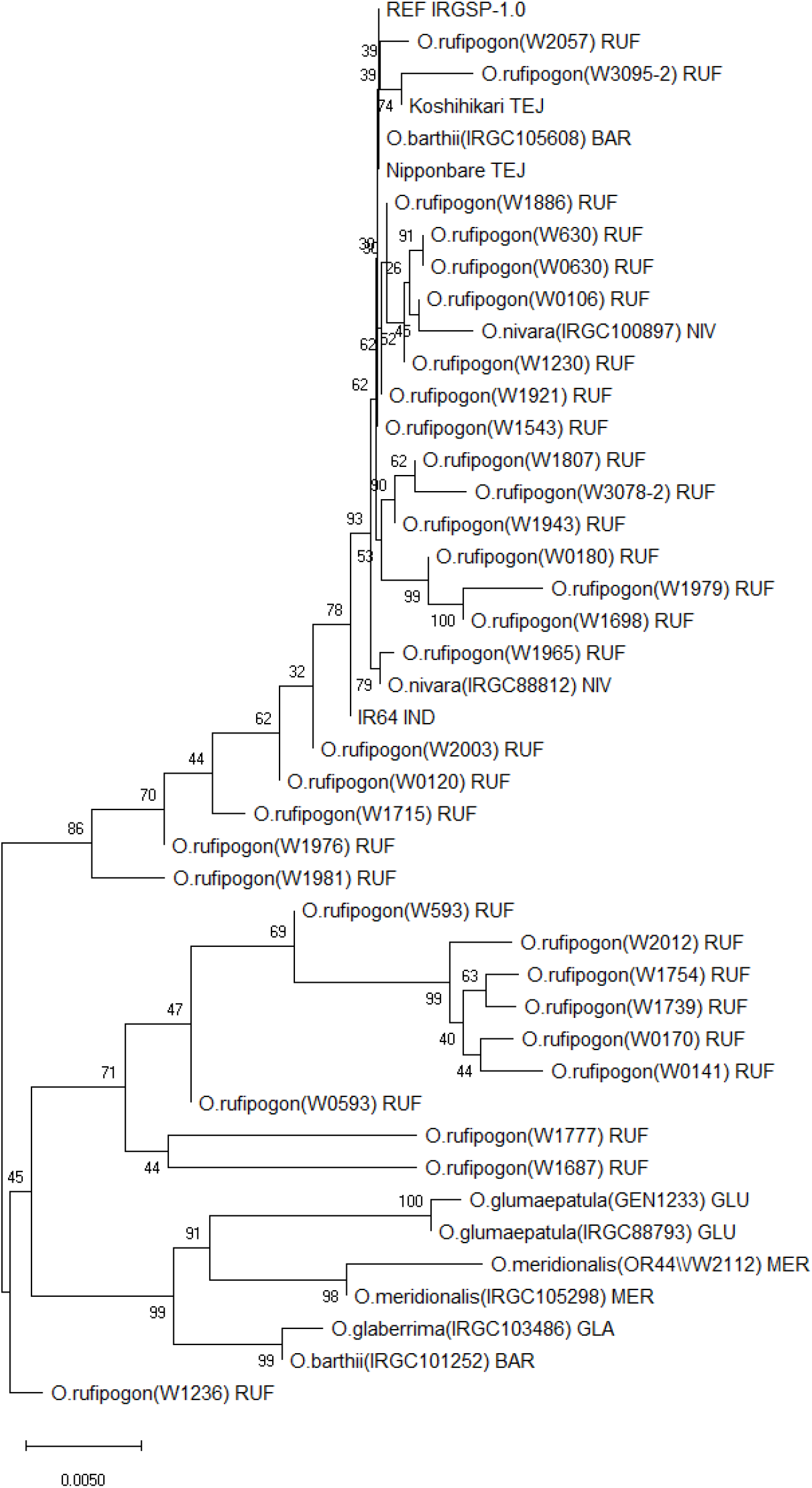
Phylogenetic analysis of DECUSSATE gene in wild relatives of rice

## Discussion

The first findings of this study are qDTY12.1 region (14-18Mbp) in rice has evolved differently and contributed to both upland adaptation and drought tolerance in rice. The differential grouping of Vandana (upland cultivar but drought susceptible at reproductive stage) and N22 (drought tolerant) in phylogeny for different genomic regions within qDTY12.1, differential fixation of DECUSSATE alleles in rice ecotypes, and diversity of OsDEC gene sequences in wild rice relatives substantiates the evolutionary significance of genomic blocks within qDTY12.1. In agreement with our findings, OsNCED2^T^ allele was found only in upland cultivars (Lyu et al, 2013) and enhanced ABA levels and lateral root density contributing to aerobic adaptation of rice (Hu et al, 2020). Additionally, a comprehensive GWAS analysis using 3.3 million SNPs also identified several significant root length associated SNPs in the entire 12^th^ chromosome specifically in indica than japonica ecotypes (Zhao et al, 2018). In rice, 11^th^ and 12^th^ chromosome was found to be evolved and associated with richness of disease resistance R genes (Rice chromosome sequencing consortium, 2005). Similarly, we presume that the 12^th^ chromosome may also be differentially evolved and associated for ecotypes diversification and drought tolerance in rice. However, rather than gene duplications which are responsible for multiple homologs of R genes in 11 and 12^th^ chromosome, evolutionary mechanisms might be different in 12^th^ chromosome especially within qDTY12.1 genomic regions which are associated with combinations of drought tolerance, root biomass and aerobic adaptations.

The second findings from this study is the possible ‘omnigenic effect’ of OsNCED2^T^ allele with qDTY12.1 in rice. The omnigenic model was initially proposed for disease related traits in humans wherein strong trait associated loci along with adjoining genes and other genes spread in the genome are interconnected and regulates the phenotype of complex traits (Boyle et al, 2017). Recently, de los Reyes (2021) identified DECUSSATE gene in rice as one of the causal gene in qDTY12.1 which regulates panicle development under drought stress. Additionally, another causal gene OsNAM_12.1_ and several other candidate genes have been reported for controlling the drought tolerance mechanism in qDTY12.1 (Dixit et al, 2015). All these reports strengthen the omnigenic effect of genes in qDTY12.1 for conferring yield advantage under drought stress. Since, both the OsNCED2^T^ (Hu et al, 2020) and OsNAM_12.1_ (Dixit et al, 2015) regulates root length and grain yield, Vandana cultivar consist of favorable allele of NCED2 gene, and it is located within the genomic regions studied for the characterization of qDTY12.1, we propose that OsNCED^T^ gene functions as omnigenic gene and regulates the qDTY12.1 for the maintenance of yield under severe drought stress. The enhanced ABA levels and signaling due to OsNCED^T^ allele might be considered as omnigenic mechanism for the qDTY12.1 conferring drought tolerance. However, this assumption requires further studies and characterization.

The development of water saving rice varieties has the potential to ensure sustainable rice production in coming decades (Luo et al, 2010). In this regard, qDTY12.1 which showed consistent yield performance and contributed to 23% phenotypic variance for yield under severe drought stress conditions is highly essential for improvement of rice varieties (Mishra et al, 2013). Additionally, membrane stability QTLs has also been identified in rice for drought tolerance (Tripathy et al, 2000; Shanmugavadivel et al, 2019). Thus, a breeding model through introgression of NCED2^T^, qDTY12.1, and QTLs/genes conferring membrane stability for drought stress has been proposed for further enhancement of genetic gain for yield under severe drought stress in rice (Fig. 7). The proposed model incorporates the physiological processes namely rapid response, yield maintenance, and membrane stability under severe drought stress conditions.

**Fig. 7.**
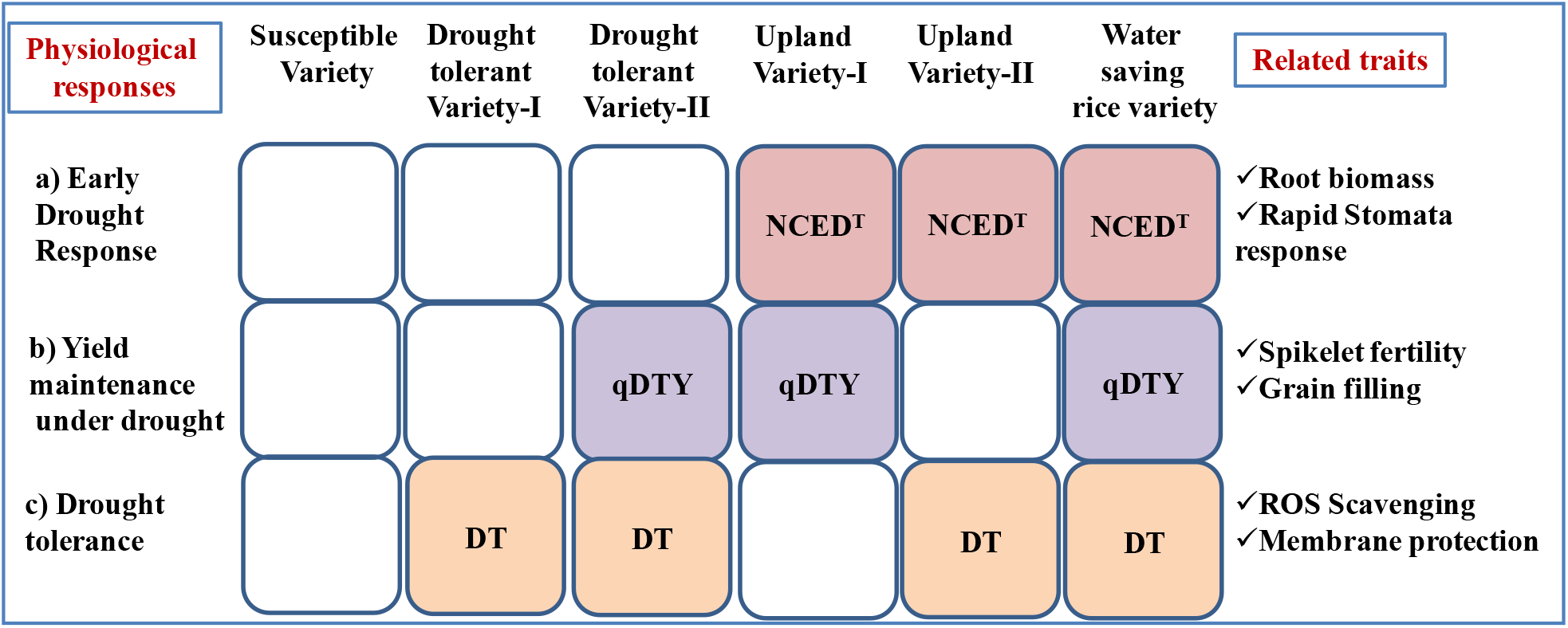
Schematic representation for enhancing genetic gain for yield under severe drought stress in rice. Breeding models for the development of water saving rice varieties in the present model comprises of combination of early drought response (pink color), yield maintenance (purple color) and drought tolerance (brown color) conferred by NCED^T^, qDTY12.1, and drought tolerant QTLs. White color indicates absence of above mentioned traits in susceptible varieties. Water saving varieties comprises of all three physiological responses under severe drought stress conditions. Upland or drought tolerant varieties comprises of any two of above mentioned physiological responses. The related traits for these three physiological responses are given in right side of the schematic diagram.

